# Engineered AAV9 as *in vivo* gene delivery platform for the selective transduction of TME cell subsets

**DOI:** 10.1101/2025.06.11.658408

**Authors:** M. B. Demircan, F. Straßheimer, L. J. Zinser, P. Elleringmann, F. John, F. B. Thalheimer, J. P. Steinbach, T. Oellerich, M. C. Burger, C. J. Buchholz

## Abstract

Precise *in vivo* gene delivery to specific cell types remains a significant challenge in gene therapy, particularly for cancer immunotherapy applications. Here, we rationally engineered AAV9 to become a modular, receptor-targeted vector for selective *in vivo* gene delivery. We first identified the N272A and W503A mutations as effective in ablating the native tropism of AAV9. Subsequently, designed ankyrin repeat proteins (DARPins) were inserted into the GH2/3 capsid loop, redirecting vector specificity towards defined cellular receptors without compromising capsid integrity or yield. As a proof of concept, HER2-targeted DART-AAV9 vectors demonstrated highly selective transduction of HER2-positive tumor cells *in vitro* and *in vivo*, in both, subcutaneous and orthotopic glioblastoma models, with negligible transduction of off-target organs including liver, heart, and kidney. When equipped with immunomodulatory genes (anti–PD-1 or IL-2) HER2-DART-AAV9 mediated secretion of functional therapeutic proteins from transduced tumor cells. Additionally, our modular platform facilitated rapid generation of CD8-targeted DART-AAV9 vectors, enabling selective transduction of human CD8^+^ T cells. Importantly, the engineered vectors exhibited favorable resistance to neutralization by human serum and retained their specificity and potency in human blood, underscoring their potential for clinical translation. Together, these findings establish DART-AAV9 as a versatile, precise, and clinically promising gene delivery platform for cancer immunotherapy.

## Introduction

Effective localized delivery remains a significant challenge in cancer immunotherapy, as systemic administration of immunomodulating agents can lead to off-target effects and dose-limiting toxicities. To address these limitations, various approaches have been explored, including nanoparticles, lipid nanoparticles (LNPs), microneedle-based systems, hydrogels, combined delivery platforms, and implantable devices designed for localized therapeutic release, all demonstrating promising activities in preclinical tumor models^1, 2^. While these studies emphasize the potential therapeutic benefit of localized immune modulation, these technologies still face notable limitations, primarily the need for direct intratumoral or peritumoral administration. Such localized interventions are viable mainly for superficial or easily accessible lesions and remain impractical for deep-seated, multifocal, or metastatic tumors, leaving disseminated tumor cells untreated^3^. Therefore, there is an urgent need for delivery systems capable of selectively targeting therapeutic genes specifically to the tumor microenvironment (TME) via systemic administration. Among available strategies, vector engineering for tumor-targeted delivery of genetic payloads harbors significant promise for achieving the required level of selectivity^4–7^.

Among viral vectors, adeno-associated virus (AAV) has rapidly become integral to contemporary therapeutic strategies due to its low immunogenicity, favorable safety profile, and robust transgene expression. Over 200 clinical trials and several market approvals have validated AAV’s therapeutic potential and safety. Notably, AAV-based therapies such as Luxturna for inherited retinal disease (Leber congenital amaurosis) and Zolgensma for spinal muscular atrophy have demonstrated transformative clinical outcomes, effectively addressing previously untreatable genetic conditions^8–10^. Notably, Zolgensma utilizes AAV9, a naturally occurring serotype, known for its potent tissue penetrance and unique capability to cross the blood-brain barrier (BBB). However, AAV9’s potency is accompanied by substantial off-target effects, particularly a 300–1,000-fold higher transduction rate in the liver compared to the intended central nervous system targets^11^. Despite these challenges, the clinical success achieved with AAV9 underscores the considerable potential for purpose-specific engineering of AAV9 capsids to optimize therapeutic efficacy and minimize off-target effects.

Following systemic administration, AAV particles often distribute widely and transduce non-target tissues. The broad tropism and especially liver accumulation of AAV9 and other serotypes has led to severe dose-limiting toxicities including fatal cases due to hepatotoxicity^11–13^. Effective cancer gene therapy therefore requires directing vectors to specific cell subsets within the TME. The importance of this precision is underscored by recent shifts in oncology: as the TME’s role in tumor progression became evident, cells of the TME (e.g. endothelial cells, macrophages, T lymphocytes) have emerged as crucial therapeutic targets alongside the tumor cells themselves. Consequently, an important challenge for gene delivery is achieving highly selective transduction of the respective TME cell types.

Recent advances in AAV bioengineering are beginning to overcome this challenge of achieving cell-type specificity. Over the past decade, researchers have focused AAV capsid engineering through approaches ranging from directed evolution to rational protein engineering. These efforts have yielded next-generation AAV variants with novel tropisms and improved targeting profiles^14^. Among these approaches, one promising strategy is receptor-targeted AAV design, wherein peptides or protein ligands are genetically inserted into the viral capsid to retarget the vector to alternative cellular receptors^15^. For example, insertion of a high-affinity binder at an exposed capsid loop can enable AAV to utilize a non-natural receptor for cell attachment/entry^16^. By introducing a novel targeting moiety, alongside disruption of native receptor binding sites, AAV tropism can be redirected toward specific cell types while minimizing off-target transduction^17^. Among targeting ligands, designed ankyrin repeat proteins (DARPins) have proven especially powerful. DARPins are engineered binding proteins that combine high affinity and specificity in a compact, highly-stable scaffold^18^. Incorporating DARPins into AAV2 and AAV6 capsids has enabled highly selective transduction of target cells. For instance, DARPin-targeted AAVs have been developed for neuronal and immune cell markers^19–22^. A recent study reported an AAV engineered to display a CD8-specific DARPin, yielding a vector that targets CD8^+^ cytotoxic T cells with remarkable specificity. *In vivo*, this CD8-targeted AAV transduced up to 80 % of CD8^+^ T cells after a single intravenous injection, while achieving near-complete exclusion of off-target cells and minimal liver transduction^22^. This example demonstrates how AAV capsid engineering can confer highly selective cell targeting, enabling precise *in vivo* gene delivery in complex multicellular environments.

Capitalizing on these advances and on AAV9’s distinctive attributes of broad tissue tropism, deep parenchymal penetration, and efficient BBB passage, this study introduces a modular receptor-targeted AAV9 vector platform tailored for cancer immunotherapy. We engineered two distinct but complementary vectors: HER2-AAV9 targeting HER2-positive tumor cells and CD8-AAV9 targeting cytotoxic T lymphocytes (CTLs). HER2-AAV9 aims for localized gene delivery directly to tumor cells, exploiting HER2 receptor overexpression common in various malignancies, while CD8-AAV9 selectively modulates CD8^+^ T cells, key effectors of anti-tumor immunity. This dual-vector strategy allows precise modulation of both tumor cells and immune effectors, enhancing therapeutic efficacy and specificity. Additionally, the vectors’ modular nature permits rapid retargeting to alternative cellular receptors or payloads, underscoring their versatility and potential broad applicability. Thus, the HER2-AAV9 and CD8-AAV9 platform represents a significant advancement toward effective, targeted, and systemic gene therapy for cancer.

## Results

Our strategy to generate DART-AAV9 from the unmodified AAV9 capsid involved two key modifications: (1) disrupting the natural receptor binding (terminal galactose) through “blinding” mutations, and (2) inserting a target-specific DARPin to refunctionalize the capsid for target cell binding and entry. Various AAV9 capsid mutations have previously been suggested to effectively abolish galactose binding, its natural primary attachment factor^23, 24^. To identify an optimal blinding mutation, we introduced four sets of distinct mutations L380A/T381A, N562A/E563A, N704A/Y705A or N272A/W503A into the AAV9 capsid and evaluated their impact on transduction efficiency on HUH7, SH-SY5 and GL261-HER2 cell lines relative to unmodified AAV9. All tested mutations substantially decreased transduction across all cell lines, the L380A/T381A mutation, however, exhibited significant residual activity on HUH7 and GL261-HER2 cells (Fig. S1A).

Next, we assessed whether DARPin insertion could restore targeted transduction in these blinded AAV9 capsids. For initial screening, we generated chimeric AAV variants carrying the HER2-specific DARPin 9.29 inserted into VP1 of AAV2, combined with VP1, VP2, and VP3 from AAV9 harboring one of the blinding mutations sets: N562A/E563A, N704A/Y705A or N272A/W503A. Among these constructs, both HER2-AAV2/9^N704A/Y705A^ and HER2-AAV2/9^N272A/W503A^ showed enhanced transduction into HER2-positive GL261-HER2 cells, while HER2-negative cells were spared (Fig. S1B). Based on these findings, we selected the N704A/Y705A and N272A/W503A mutations for subsequent testing in a full AAV9 background.

Following this initial screening with AAV2-VP1-DARPin constructs, which yielded functional chimeric capsids, we generated fully AAV9-based receptor-targeted capsids. To achieve this, we employed a dual-plasmid approach: one plasmid encoding VP1 with blinding mutations and insertion of the coding sequence of the DARPin into the GH2/3 loop and the second plasmid encoding VP2,3 with the same blinding mutations. Additionally, DARPin-inserted AAV9 without blinding mutations was generated (HER2-AAV9^noblind^). Thus, HER2 DARPins were inserted into the GH2/3 loop of AAV9 VP1 either alone or in combination with the N704A/Y705A or N272A/W503A blinding mutations. These VP1 variants were co-expressed with the corresponding VP2/3 plasmids, either lacking or containing the corresponding mutations. Upon transduction of HER2-positive glioma cells (GL261-HER2 and LN-319), HER2-AAV9^noblind^ showed enhanced transduction compared to unmodified AAV9. However, although transduction in HER2-negative cells (GL261 and SH-SY) was reduced, substantial residual activity persisted. Among the blinded variants, HER2-AAV9^N272A/W503A^ demonstrated increased transduction of HER2-positive cells over unmodified AAV9 while showing negligible transduction of HER2-negative cells (Fig. S2A, B). However, the overall transduction efficiency of HER2-AAV9^N272A/W503A^ variant was lower than that of HER2-AAV9^noblind^. We hypothesized that mutating two critical regions of the capsid (N272A and W503A) may have compromised overall activity. To address this, we proceeded to test single-mutation variants. Both, HER2-AAV9^N272A^ and HER2-AAV9^W503A^, exhibited enhanced transduction in HER2-positive cells compared to HER2-AAV9^N272A/W503A^ without increasing transduction in HER2-negative cells (Fig. S2B). Based on these results, we selected the single W503A mutation as the preferred blinding mutation for the generation of DART-AAV9 in this study. From here on, the corresponding AAV variant will be designated as HER2-AAV9.

Next, we performed a series of biochemical and biophysical characterizations of HER2-AAV9 vector particles (Fig. 1A), using iodixanol-purified vector stocks. Western blot analysis confirmed successful DARPin incorporation into both HER2-AAV9 variants, as evidenced by a shifted VP1 band relative to that of unmodified AAV9. Notably, an additional band migrating at the expected molecular weight of a VP3-DARPin fusion protein suggested residual VP3 expression despite the disruption of the major splice acceptor site downstream of the VP1 initiation codon (Fig. 1B). DARPin displaying AAV9 capsids retained a high thermal stability, with only a modest decrease relative to unmodified AAV9 (Fig. 1C). Importantly, several production batches of the various DARPin-displaying AAV9 capsids revealed high reproducibility in production, with vector titers in the same range as those of unmodified AAV9 (Table S1). In functional assays, HER2-AAV9^noblind^ exhibited dose-dependent transduction of both HER2-positive and HER2-negative cells, whereas HER2-AAV9 displayed high-level of transduction exclusively in HER2-positive cells, remaining completely inactive on HER2-negative cells (Fig. 1D, E). Finally, competition assays indicated that HER2-AAV9 transduction were reduced significantly in the presence of a HER2-specific antibody, further supporting the specificity of the engineered DARPin-capsid interaction (Fig. S3).

**Figure 1:**
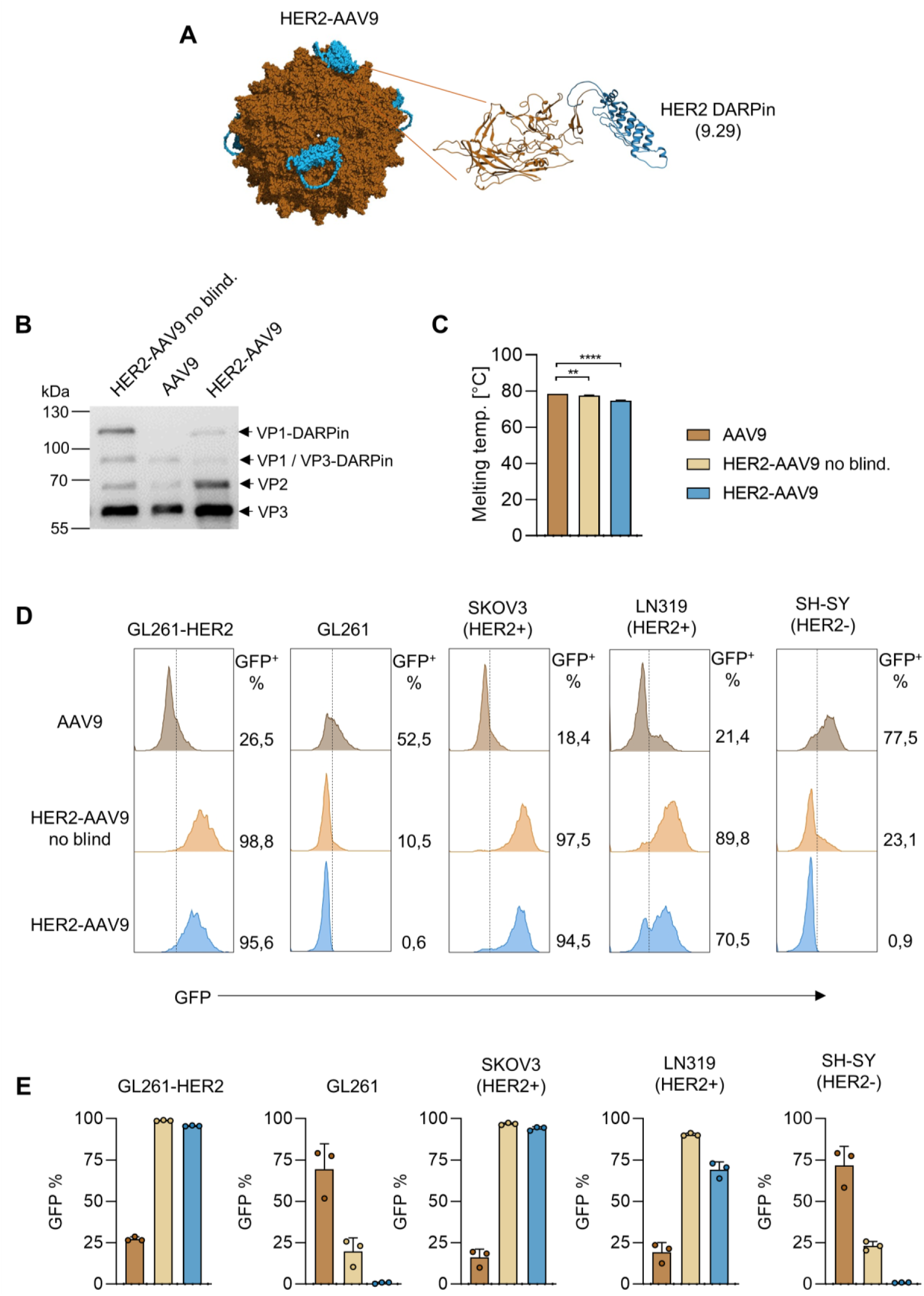
Design and characterization of HER2-AAV9 vectors. **A)** Left: AAV9 capsid based on the structure of AAV9-VP3 (PDB: 3UX1). AlphaFold 3 structure predictions of HER2-AAV9 were aligned with single copies of VP3 in the capsid. Right: AAV9-VP3-9.29 proteins with N-terminal (G_4_S)_5_ linker (HER2-AAV9). 9.29 DARPin and linker are shown in blue and VP3 in brown. **B)** Western blot analysis of the indicated HER2-AAV9 vectors. **C)** Differential scanning fluorimetry of the indicated AAV stocks. 1 x 10^10^ vg/µl of each sample were exposed to increasing temperatures (1.5 °C/min) from 30 °C to 95 °C (n = 3). Statistical analyses were performed using unpaired t-test. Bars represent means, error bars represent SD, p-value ** < 0.01 and **** < 0.0001. **D, E)** GFP reporter gene transfer mediated by the indicated vectors on HER2+ (GL261-HER2, SKOV3, LN-319) and HER2- (GL261, SH-SY) cells. Cells were incubated with the indicated AAVs at 1 x 10^5^ vg/cell. **(D)** Representative flow cytometry histograms showing GFP signal analysis at 3 days post-transduction are shown. **(E)** Transduction efficiency was determined as the percentage of GFP-positive cells within all live cells (n = 3). Bars represent means, error bars represent SD.

To evaluate the potential of the newly developed AAV9 variants, we tested a range of HER2-targeted AAV vectors, including a previously reported first-generation HER2-AAV2 vector featuring a DARPin insertion at the N-terminus of VP2 ^17^ (HER2-AAV2^N-term^), a second-generation HER2-AAV2 vector with DARPin insertions in the GH2/3 loop (HER2-AAV2^Loop^), and the corresponding HER2-AAV9 variants. All capsid versions, each encoding luciferase, were initially assessed for transduction efficiency on HER2-positive cell lines (SKOV3, LN-319 and GL261-HER2). While the first-generation HER2-AAV2^N-term^ conferred either an approximately 12-fold increase or a 3-fold decrease in transduction in various cell lines relative to unmodified AAV9, the second-generation variants achieved substantially higher transduction efficiencies in all tested cell lines. Specifically, HER2-AAV2^Loop^ insertions achieved up to 271-fold increase, HER2-AAV9^noblind^ up to 531-fold, and HER2-AAV9 up to 474-fold (Fig. S4A-C).

Next, we evaluated the performance of these capsids *in vivo* using nude mice bearing subcutaneous HER2-positive SKOV3 xenografts. Seven days following systemic AAV injection, mice were imaged for luciferase activity and then sacrificed to collect tumors and relevant organs for further analysis. Imaging revealed that all targeted AAVs effectively minimized off-target transduction, while exhibiting varying levels of on-target transduction in HER2-positive tumors (Fig. 2A and Fig. S5A). To obtain quantitative data, tumors and primary AAV9-targeted organs were lysed, and luciferase activity was measured via luciferase assay. Compared with unmodified AAV9, HER2-AAV9^noblind^ and HER2-AAV9 exhibited 6-fold and 4-fold increases in tumor-specific luciferase signals, respectively. In contrast, HER2-AAV2^N-term^ and HER2-AAV2^Loop^ were less active in transducing the tumor (Fig. 2B). Importantly, HER2-AAV9, HER2-AAV2^N-term^, and HER2-AAV2^Loop^ variants were all completely detargeted from liver with signals indistinguishable from PBS controls (Fig. 2C). Moreover, transduction in other highly permissive organs such as heart and kidney were also dramatically reduced (Fig. S5A-C). Among all tested vector variants, HER2-AAV9 emerged as particularly effective *in vivo*,

**Figure 2:**
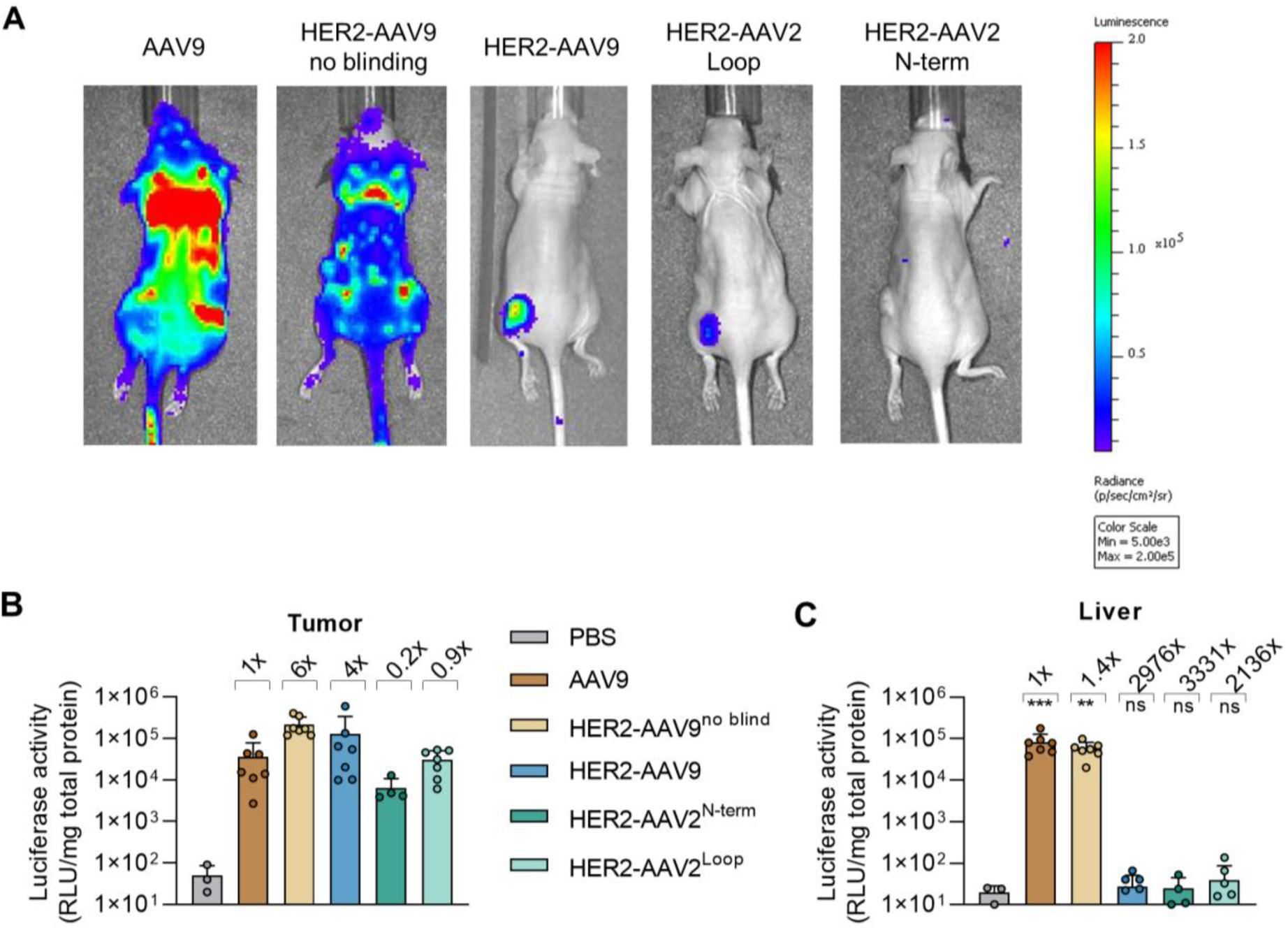
HER2-AAV9 enables tumor-specific gene delivery *in vivo*. *In vivo* transduction in a subcutaneous tumor model. Nude mice were engrafted with SKOV3 cells in the left flank and systemically injected via the tail vein with each luciferase-encoding AAV vectors (2 x 10^11^ vg). **(A)** Representative whole-body IVIS images at day 7 post-injection are shown. Following euthanasia, tumors and organs were harvested, lysed in luciferase lysis buffer, and analyzed to determine transduction efficiency in tumors **(B)** and livers **(C)**. Each dot represents an individual mouse (n = 7 except of n = 3 for PBS control and n = 4 for HER2-AAV2^N-term^). The fold increase **(B)** or decrease **(C)** in transduction relative to AAV9 is indicated above each bar. Statistical analyses were performed using unpaired t test. Bars represent mean values; error bars represent SD. p-value ** < 0.01, ∗∗∗∗p < 0.0001.

To further capitalize on the unique ability of AAV9 to cross the blood–brain barrier (BBB), an especially valuable feature for targeting brain tumors, we next investigated the performance of HER2-AAV9 in an orthotopic glioblastoma model (Fig. 3A). Mice bearing intracranial HER2-positive GL261 tumors received intratumoral injections of the indicated AAV vectors. Both unmodified AAV9 and HER2-AAV9^noblind^ mediated transduction not only within the tumor but also in the contralateral (non-tumor) hemisphere, indicating off-target transduction of normal brain tissue. In contrast, HER2-AAV9 and HER2-AAV2^Loop^ demonstrated striking selectivity, achieving robust gene transfer specifically within HER2-expressing glioblastoma tumors and exhibiting minimal transduction in the contralateral hemisphere (Fig. 3B). Additionally, only unmodified AAV9 displayed detectable off-target transduction in the liver, highlighting the detargeting achieved with engineered capsids (Fig. 3C).

**Figure 3:**
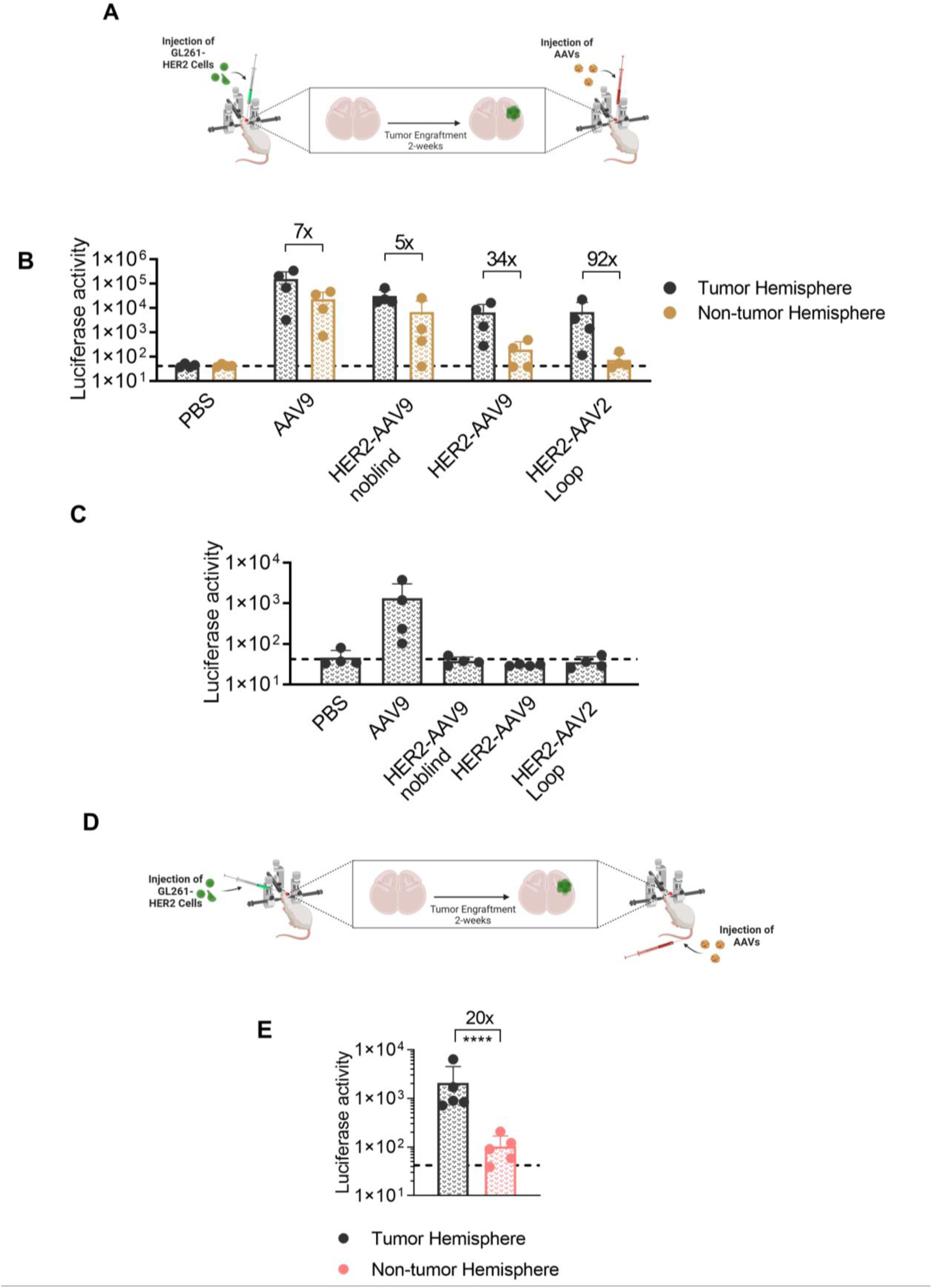
HER2-AAV9 enables tumor-specific gene delivery in brain orthotopic tumor models. **A-C)** Syngeneic C57BL/6 mice were engrafted with GL261-HER2 cells in the right striatum and intratumorally injected with the indicated luciferase-encoding AAV vectors. **(A)** Experimental setup. Following euthanasia, organs were harvested, lysed in luciferase lysis buffer, and analyzed to determine transduction efficiency in the right tumor hemisphere and left non-tumor hemisphere **(B)** and livers **(C)**. **(D-E)** Syngeneic C57BL/6 mice were engrafted with GL261-HER2 cells in the right striatum and systemically injected via the tail vein with the luciferase-encoding HER2-AAV9 vector. **(D)** Experimental setup. **(E)** Following euthanasia, organs were harvested, lysed in luciferase lysis buffer, and analyzed to determine transduction efficiency in the right tumor hemisphere and left non-tumor hemisphere). Each dot represents an individual mouse (n = 4-5). The fold increase in transduction relative to tumor versus non-tumor hemisphere is indicated above bars. Statistical analyses were performed using 2-tailed ratio paired t test. Bars represent mean values; error bars represent SD. p-value ∗∗∗∗p < 0.0001. exhibiting both potent and highly selective tumor transduction with minimal detectable off-target activity.

We further assessed systemic administration of HER2-AAV9 in the same orthotopic glioblastoma model (Fig. 3D). Notably, a single intravenous injection of HER2-AAV9 resulted in selective transduction within intracranial HER2-positive tumors, with background-level transgene expression in normal brain tissue similar to PBS-treated controls (Fig. 3E).

To explore the potential of engineered HER2-AAV9 for targeted immunotherapy, we packaged the coding sequences for the immune checkpoint inhibitor mouse anti-PD1, the human cytokine IL2, and a CD122-biased mutant of IL2 (IL2_A4; designed to reduce Treg activation)^25, 26^, into HER2-AAV9. Transduction of GL261-HER2 and SKOV3 cells with HER2-AAV9^anti-PD1^ vectors resulted in the secretion of anti-PD1 protein into the culture supernatant, which, upon incubation with PD1-expressing M58ab mouse cells demonstrated binding to PD1 on the cell surface (Fig. 4A, B). Similarly, the supernatant from cells transduced with HER2-AAV9 IL2 and HER2-AAV9 IL2_A4 vectors significantly enhanced the proliferation of an IL2-dependent NK cell line (KHYG-1), indicating successful transduction and secretion of functional IL2 by HER2-positive cells (Fig. 4C).

**Figure 4:**
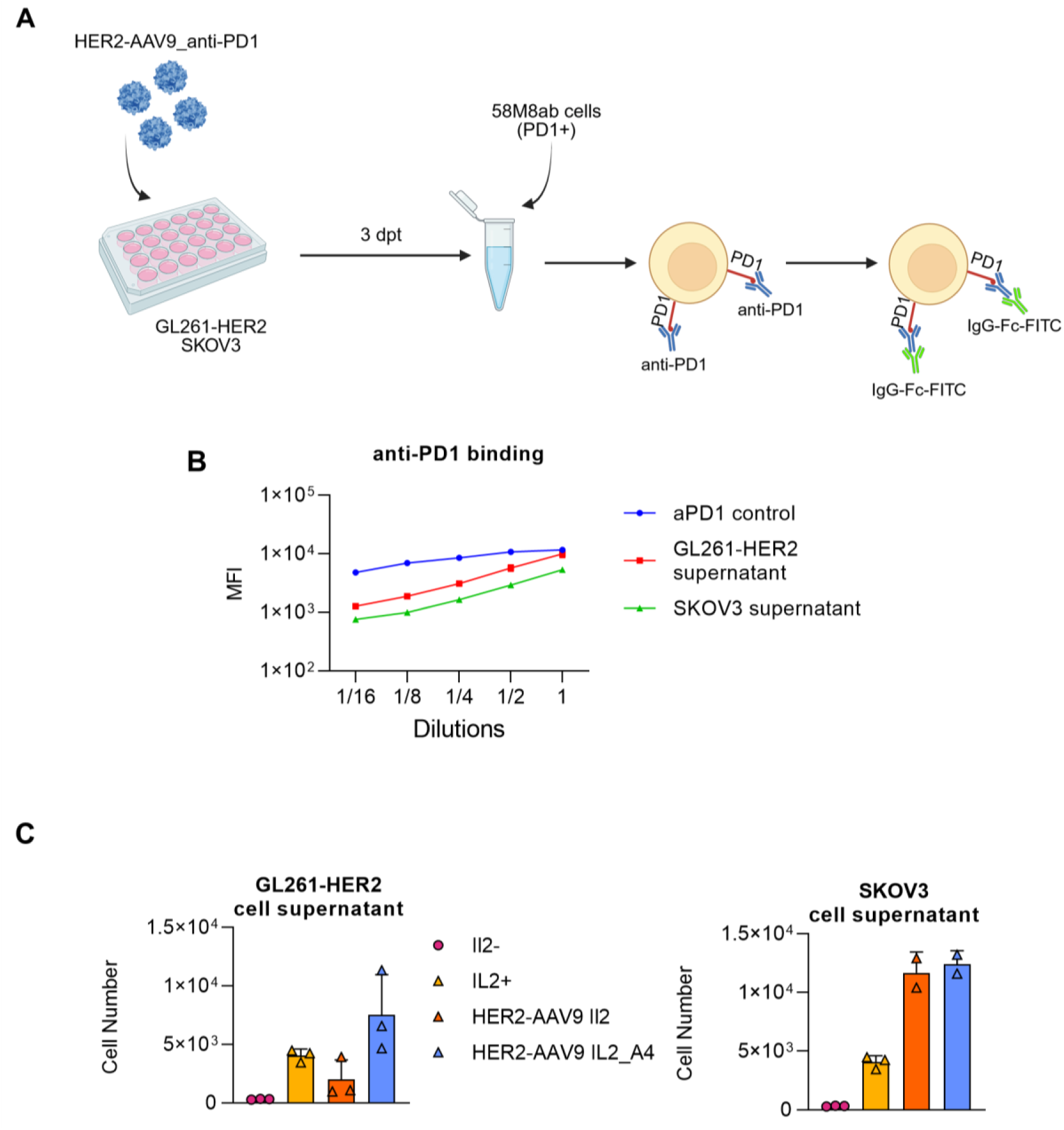
HER2-AAV9 mediated delivery of immune-modulators to HER2+ tumor cells. **A)** Experimental setup: HER2+ cells were transduced with HER2-AAV9 vectors (1 x 10^5^ vg/cell) encoding a mouse anti–PD1 antibody as transgene. On day 3 post-transduction, the culture supernatant was collected to measure anti–PD1 secretion. The collected supernatant was then incubated with 58M8ab cells expressing mouse PD1, and anti– PD1 binding was detected by flow cytometry using a FITC-conjugated secondary antibody. 2 µg/ml anti–PD1 solution served as a positive control. **B)** Mean fluorescence intensities (MFIs) at different supernatant dilutions are shown (n = 3). **C)** HER2+ cells were transduced with HER2-AAV9 vectors (1 x 10^5^ vg/cell) encoding IL2 or IL2_A4. On day 3 post-transduction, culture supernatants were collected to assess IL2 functionality. Each supernatant was diluted 1:1 with culture medium and used to culture the IL2-dependent KHYG-1 cell line. 2 µg/ml IL2 solution served as a positive control. Cell counts on day 5 are shown (n = 3). Bars represent mean values; error bars represent the SD.

Our rational AAV capsid engineering approach enables rapid and flexible modification of target specificity, significantly broadening their applicability across various therapeutic settings. Beyond targeting tumor cells directly within the TME, another promising complementary strategy in cancer immunotherapy involves direct modulation of cytotoxic T lymphocytes *in vivo*. Toward this goal, we sought to generate a CD8-targeted AAV9 vector capable of selectively transducing and modifying CD8-positive T cells. The hCD8-specific DARPin 53F6 ^27^ was chosen as a high-affinity targeting ligand for human cytotoxic T cells. For display on the surface of AAV9 vector particles, the coding sequence of the previously described HER2 DARPin 9.29 was exchanged for that of CD8 DARPin 53F6 (Fig. 5A). When tested on human donor peripheral blood mononuclear cells (PBMCs), hCD8-AAV9 showed slightly lower overall transduction efficiency compared to the previously developed hCD8-AAV2 vector. However, hCD8-AAV9 substantially improved CD8+ T-cell transduction, increasing the average rate from 8.5 % with unmodified AAV9 to 78 % (Figures 5B, C). Notably, the selectivity of hCD8-AAV9 for CD8^+^ cells approached nearly complete specificity at 99.1 %, surpassing the selectivity observed with hCD8-AAV2 (94.2 %), while unmodified AAV9 demonstrated non-selective transduction with only 55.7 % specificity (Figure 5D).

**Figure 5:**
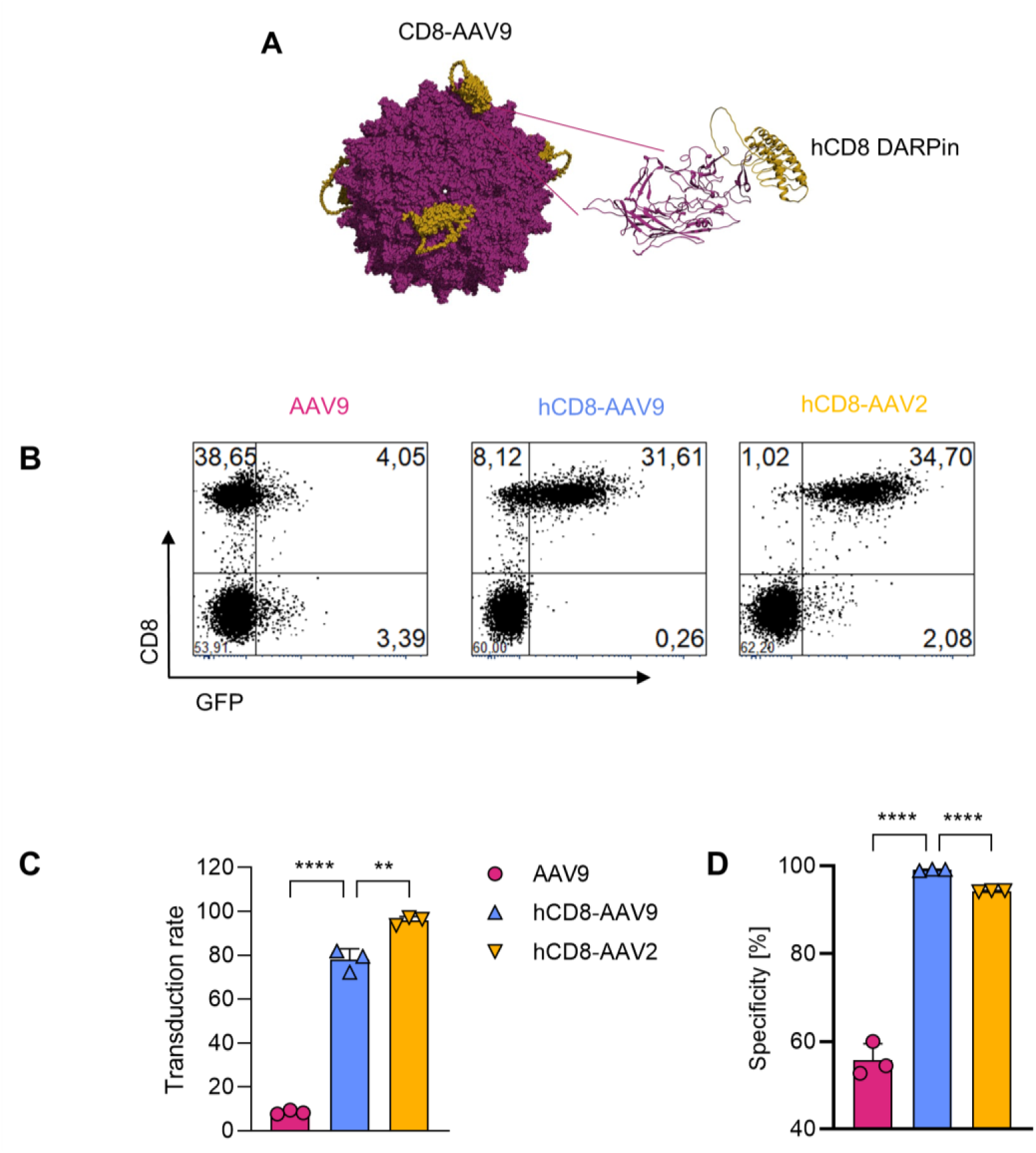
CD8-AAV9 provides selective gene transfer into human PBMCs. **(A)** Left: AAV9 capsid based on the structure of AAV9-VP3 (PDB: 3UX1). AlphaFold 3 structure predictions of CD8-AAV9 were aligned with single copies of VP3 in the capsid. Right: AAV9-VP3--53F6 proteins with N-terminal (G_4_S)_5_ linker (CD8-AAV9). 53F6 DARPin and linker are shown in yellow and VP3 in magenta. **(B–D)** Transduction of activated human PBMCs isolated from three different donors (n = 3) using the indicated AAV vectors encoding GFP at 5 × 10^5^ vg/cell. **(B)** Representative flow cytometry plots are shown. **(C)** Transduction efficiency was quantified as the percentage of GFP-positive cells within the CD3+/CD8+ T-cell population. **(D)** Transduction specificity was determined as the percentage of CD8+ T cells among all GFP-positive cells. Statistical analyses were performed using unpaired t-test. Bars represent means, error bars represent SD, p-value ** < 0.01 and **** < 0.0001

To assess the translational potential of our engineered AAV vector platform for clinical application, we first evaluated their susceptibility to neutralization by inhibitory factors present in human serum. HER2-AAV9 and unmodified AAV9 vectors were incubated with serial dilutions of pooled human serum, followed by assessment of transduction efficiency. Interestingly, HER2-AAV9 was significantly less sensitive towards neutralization at a low serum concentration than unmodified AAV9, suggesting that the capsid modification decreased the vulnerability to inhibitory serum components and neutralizing antibodies (Fig. 6A).

**Figure 6:**
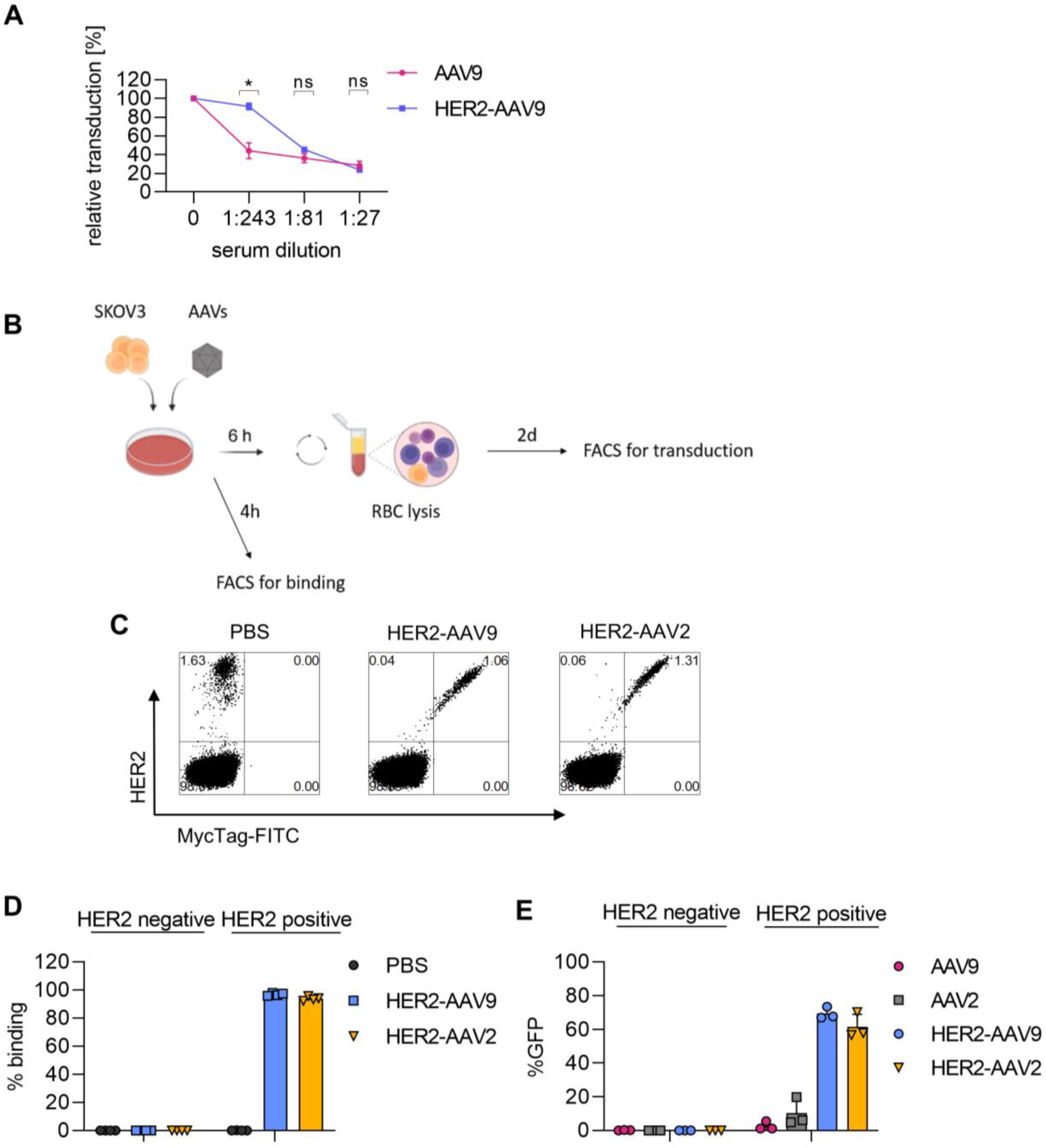
Her2-AAV9 targets tumor cells in human blood. **(A)** Effect of human serum pre-incubation on transduction efficiency of AAV9 and HER2-AAV9 vectors. Vectors encoding GFP were pre-incubated with three-fold serial dilutions of pooled human off-the-clot serum and subsequently added to target cells for transduction. GFP expression was measured by flow cytometry at 3 days post-transduction. Percent inhibition of transduction was calculated by normalizing GFP expression levels to those obtained from vectors incubated without serum. Statistical analyses were performed using multiple t-test ∗p < 0.05. **(B)** Experimental setup for binding and transduction assays in human blood. SKOV3 cells were mixed with human blood, and the cell-blood mixture was incubated with AAV vectors at a concentration of 2.5 × 10^10^ vg/ml blood. **(C-D)** For the binding assay, cells were incubated with vector particles for 4 hours, followed by red blood cell (RBC) lysis and analysis by flow cytometry. **(C)** Representative flow cytometry plots are shown. **(D)** % binding was determined as the percentage of myctag-FITC positive cells in HER2 negative and positive cells. **(E)** For the transduction assay, cells were incubated for 6 hours, washed to remove unbound AAV particles, subjected to RBC lysis, and then cultured for an additional 3 days prior to assessing transduction levels by flow cytometry (n = 3). Bars represent means, error bars represent SD.

Next, to mimic a physiologically relevant environment, HER2^+^ SKOV3 cells were mixed into human blood and treated with the indicated AAV vectors. Binding and transduction assays were subsequently performed (Fig. 6B). Despite the low abundance (1-2 %) of HER2^+^ cells within the whole blood cell mixture, HER2-AAV9 specifically bound to the target population with minimal off-target interaction (Fig. 6C, D). Consistent with the binding data, both HER2-AAV9 and HER2-AAV2 mediated robust and selective transduction of HER2^+^ cells in human blood. In contrast, unmodified AAV9 and even AAV2, which typically exhibits high *in vitro* activity, failed to achieve meaningful transduction under these conditions (Fig. 6E). Importantly, in a parallel experiment, CD8-AAV9 vectors also exhibited highly selective binding to CD8^+^ T cells in human blood, with minimal interactions with non-target cell populations (Fig. S6). These findings highlight the specificity and functionality of targeted AAV vectors in the complex environment of human blood, supporting their potential for *in vivo* clinical applications.

## Discussion

We present a modular platform for receptor-targeted AAV9 vectors (DART-AAV9), enabling cell-type–specific *in vivo* gene delivery. AAV9 offers unique advantages due to its broad native tropism and inherent ability to efficiently traverse the blood–brain barrier (BBB). Leveraging AAV9 for selective gene delivery required overcoming two critical challenges: (1) ablating its natural glycan receptor interactions (“blinding” its intrinsic tropism), and (2) incorporating DARPins as targeting ligands into the capsid without compromising capsid integrity or functionality. Previous structural analyses have identified key residues, notably N272 and W503, on the AAV9 capsid surface as essential for terminal galactose binding^23, 24^. Substitution of these residues individually with alanine (N272A or W503A) effectively disrupted native receptor interactions, significantly reducing off-target transduction in non-target cells. Unlike AAV2, which requires a double mutation (R585A and R588A) to achieve similar effects^28^, we found that the AAV9 double mutant (N272A/W503A) impaired gene transfer activity, while the single mutations were sufficient to eliminate receptor binding while better preserving capsid integrity and functionality. Following established precedents from AAV2 and AAV6 engineering, we chose the surface-exposed GH2/GH3 loop within variable region IV (VR-IV) for DARPin insertion^22^. The resulting DART-AAV9 vectors exhibited intact capsid structures and maintained high thermal stability. Importantly, DART-AAV9 vectors were consistently produced at titers comparable to unmodified AAV9, which was documented for both, HER2-AAV9 and CD8-AAV9.

Retargeting AAV9 to the HER2/neu receptor yielded highly selective and efficient gene delivery to HER2+ tumor cells. HER2-AAV9 markedly outperformed earlier-generation HER2-targeted AAV vectors built on AAV2. In the first-generation design, a HER2-binding DARPin was fused to the N-terminus of the AAV2 VP2 capsid protein, and mutations were introduced to ablate heparan sulfate binding^17^. Although this first-generation HER2-AAV2 vector demonstrated HER2-dependent transduction *in vitro* and delivered genetically encoded checkpoint-inhibitors selectively to HER2-positive tumors *in vivo*, its broader translational potential will greatly benefit from enhanced gene transfer activity and improved production yields^29^. Crucially, the new variants HER2-AAV9 and HER2-AAV2^Loop^ exhibited robust capsid assembly and consistently yielded high titers, comparable to those of unmodified AAVs, underscoring their compatibility with scalable manufacturing. In comparative studies using a subcutaneous tumor model, the second-generation HER2-AAV2^Loop^ vector achieved approximately five-fold greater *in vivo* transduction efficiency than the first-generation HER2-AAV2^N-term^ vector. Importantly, the HER2-AAV9 variant further enhanced transduction efficiency, surpassing HER2-AAV2^Loop^ by approximately four-fold. Besides reporter genes HER2-AAV9 delivered immunomodulatory transgenes to tumor cells including the anti–PD-1 checkpoint inhibitor and cytokines. The superior performance observed with HER2-AAV9 likely results from intrinsic advantages of the AAV9 capsid, including increased stability under physiological conditions, prolonged circulation times prior to blood clearance, and improved tissue penetration relative to AAV2^30–32^.

Despite their substantially enhanced transduction of target cells, both HER2-AAV9 and HER2-AAV2^Loop^ exhibited nearly complete detargeting from off-target tissues, including liver, heart, and kidney, demonstrating excellent selectivity and favorable safety profiles. Furthermore, the inherent biodistribution properties of AAV9 are particularly advantageous for systemic cancer therapy, as AAV9 uniquely exhibits efficient traversal of challenging physiological barriers, notably the blood–brain barrier (BBB)^33^. Indeed, evidence for HER2-AAV9 to pass the BBB and then target the tumor tissue was obtained in an orthotopic glioblastoma model. While these results are promising, the extent of reporter gene delivery will still need optimization to match that achieved by intracranial vector administration.

The platform character of our approach was demonstrated by targeting AAV9 to CD8^+^ T lymphocytes by simply replacing the HER2-specific DARPin with a CD8-specific DARPin. The resulting CD8-AAV9 vector efficiently and selectively bound primary human CD8^+^ T cells, mediating highly specific gene transfer *in vitro*. Our previous work demonstrated robust and selective CD8^+^ T-cell transduction using CD8-targeted AAV2 and AAV6 vectors after systemic administration^22^. Importantly, the newly developed CD8-AAV9 vector achieved even higher selectivity than CD8-AAV2.

It is instructive to contrast our DARPin-based, rational receptor-targeting approach with commonly used alternative approaches such as directed evolution. A notable example is Ark313, an AAV6 variant engineered via directed evolution, which recently demonstrated highly efficient transduction of murine T cells and facilitated CRISPR-based gene editing *in vivo*^34, 35^. While Ark313 underscores the potential of targeting mouse T cells with an AAV capsid, a key limitation of this approach lies in its fixed receptor specificity, determined by the antigen used during the selection process. In this case, the capsid was binding to murine QA2, a molecule not necessarily ideal for target cell selectivity, which however resulted from the evolutionary screen. Consequently, Ark313 is functionally restricted to specific genetic backgrounds, such as the C57BL/6 mouse strain. Critically, adapting this evolved capsid to target other cell types or translating it for use in human systems would necessitate repeating the process of selection and evolution from scratch. In contrast, receptor specificity within the DART-AAV9 platform is dictated entirely by the modular DARPin ligand, enabling rational, rapid, and flexible retargeting by simply exchanging the DARPin moiety. Thus, rather than evolving entirely new capsids for each novel target, this “plug-and-play” approach leverages the robust, extensively characterized AAV9 capsid backbone, significantly accelerating the generation of targeted vectors for various cell types. This modular versatility greatly enhances the translational potential of the DART-AAV platform, opening opportunities for precise *in vivo* genetic modification of diverse immune cell subsets and clinically relevant cell populations previously out of reach for systemic gene therapy.

The DART-AAV platform, now established for the AAV9 backbone, exhibits several attributes that enhance its potential for clinical translation. To simulate clinically relevant conditions, we assessed the performance of DART-AAV9 in human blood and serum. HER2-targeted DART-AAV9 was capable for effective and selective transduction even in the presence of pooled human serum. In spiked human PBMC samples, HER2-AAV9 specifically bound to and transduced only HER2-positive tumor cells, while CD8-DART-AAV9 selectively bound CD8^+^ T cells, with no off-target binding detected in other leukocyte populations. These findings indicate that serum proteins such as immunoglobulins and complement did not impair vector specificity or induce nonspecific uptake. Notably, at lower dilution of pooled human serum, HER2-AAV9 showed reduced sensitivity to neutralization compared to unmodified AAV9. One possible explanation is that the DARPin on the capsid may sterically mask or disrupt epitopes that would otherwise be bound by neutralizing antibodies or other serum factors. These factors together with binding/uptake by non-target cells could explain why unmodified AAV9 and AAV6 were basically inactive in this setting. One of the most compelling features of DART-AAVs is modular compatibility with multiple AAV serotypes. Initially demonstrated using AAV2 and AAV6 capsids^22^, we have now extended the versatility of this approach to include AAV9, underscoring the broad applicability and modular nature of our targeting platform. This is particularly relevant in light of the high global prevalence of pre-existing neutralizing antibodies (NAbs) against AAV estimated to affect between 40 % and 80 % of the population for common serotypes^36^. The presence of NAbs can substantially reduce gene delivery efficacy and, in some cases, render patients ineligible for therapy. With the ability to graft DARPins onto diverse AAV capsids such as AAV2, AAV6, and AAV9, DART-AAVs enable serotype selection tailored to individual patients’ immunological profiles. Moreover, this serotype flexibility may also facilitate vector readministration by allowing capsid switching to circumvent anti-capsid immune responses following an initial dose—an essential capability for treating diseases that require repeated dosing^37, 38^.

In the context of cancer immunotherapy, the dual application we demonstrate, HER2-AAV9 for tumor cell transduction and CD8-AAV9 for *in vivo* T-cell modification, represents a versatile and synergistic strategy. HER2-AAV9 enables tumor cells to act as local “factories” for checkpoint inhibitors or cytokines, enhancing immune recognition and attack. Simultaneously, CD8-AAV9 can augment the anti-tumor activity of cytotoxic T lymphocytes by delivering genes directly into effector cells. Used in tandem, these vectors could orchestrate a two-pronged immune assault on the tumor.

## Material and methods

### Plasmids and molecular cloning

The AAV9 capsid plasmid pRC29 co-encoding AAV2 rep together with AAV9 cap received was modified by overlap extension PCR to insert galactose binding mutations at position L380A/T381A, N562A/E563A, N704A/Y705A or N272A/W503A. To generate relevant VP2/VP3 expression plasmids with/without galactose binding mutations at the position N704A/Y705A or N272A/W503A, VP1 start codon (ATG) was mutated to AAG in pRC29, pRC29_N704A/Y705A or pRC29_N272A/W503A plasmids. For DARPin incorporation, the coding sequence for the HER2/neu-specific DARPin 9.29 ^39^ was inserted between residues 453 and 455 of VP1 by SwaI/XcmI digestion of pRC29, pRC29_N704A/Y705A or pRC29_N272A/W503A resulting in pRC29-(G_4_S)_5_-HER2_9.29_N704A/Y705A and pRC29- (G_4_S)_5_-HER2_9.29_N272A/W503A. These VP1 expressing plasmids also contain a deletion of the splice acceptor site downstream of the VP1 translation initiation site to prevent VP2/VP3 expression. Single mutation versions pRC29-(G_4_S)_5_-HER2_9.29_N272A and pRC29-(G_4_S)_5_- HER2_9.29_W503A were derived from pRC29-(G4S)5-HER2_9.29 N272A/W503A. Exchange of the DARPin coding sequence for the CD8-specific DARPin 53F6 ^27^ was performed via SfiI/SpeI. scAAV transfer plasmids encoding the reporter gene eGFP or luciferase were previously described^17^.

### Cultivation of cell lines

HEK-293T (ATCC CRL-11268), LN-319, and GL261 cells ^29^ were cultured in high-glucose Dulbecco’s Modified Eagle Medium (DMEM; Sigma-Aldrich, D6546) supplemented with 10% fetal bovine serum (FBS; Sigma-Aldrich, F7524) and 1% L-glutamine (Sigma-Aldrich, M4892). Cells were passaged twice a week at dilution ratios ranging from 1:5 to 1:10 using 0.25% trypsin-EDTA in PBS lacking calcium and magnesium ions. SKOV3 cells ^17^ were maintained in Advanced McCoy’s 5A medium (Sigma-Aldrich, D6546) enriched with 10% FBS and 1% L-glutamine.

### Production of AAVs

AAV vector production was carried out following established protocols with modifications based on previous methods^19, 22^. Briefly, HEK-293T cells were transiently co-transfected with a combination of plasmids, including the adenoviral helper plasmid pXX6-80, packaging plasmids for VP1 and VP2/3 expression, and a transfer plasmid (pscAAV-SFFV-eGFP or pscAAV-SFFV-Luciferase_GFP). For the scaled-up generation of DART-AAVs, plasmids were used in a ratio of 10:8:8:8, consisting of pXX6-80, the transfer vector, and complementary capsid plasmids pRC22/9-VP1-KO and pRC22/9-DARPin. Three days post-transfection, both supernatants and cell lysates were harvested. Residual plasmid DNA was enzymatically digested with Benzonase® (Merck Millipore, E1014) at a final concentration of 50 U/ml for 30 minutes at 37°C. Viral particles were purified using iodixanol gradient ultracentrifugation followed by buffer exchange into PBS containing 0.01% Pluronic F-68, using 50 kDa Amicon® Ultra-4 centrifugal filter units (Merck Millipore, UFC805024). Final AAV preparations were aliquoted and stored at −80°C.

To quantify vector genome titers, AAV stocks were subjected to DNA extraction from 3 µl of vector solution using the DNeasy Blood and Tissue Kit (Qiagen, 69506). Quantitative PCR targeting the inverted terminal repeats (ITRs) was performed using specific primers and probe ^22^ with LightCycler 480 Probes Master Mix (Roche, 04707494001) on a LightCycler 480 system (Roche).

### Western blot

Western blot analysis was conducted with slight modifications from previously described protocols^19, 22^. In short, AAV vector preparations were denatured in urea buffer (200 mM Tris-HCl [pH 8.0], 5% SDS, 8 M urea, 0.1 mM EDTA, 2.5% DTT, and 0.03% bromophenol blue) by heating at 95°C for 10 minutes. The denatured proteins were resolved by SDS-PAGE and transferred onto PVDF membranes (Amersham, UK, #10600023). Membranes were blocked and then probed overnight at 4°C with a mouse monoclonal anti-AAV VP1/VP2/VP3 antibody (clone B1, Progen, #65158) at a 1:50 dilution. The following day, membranes were incubated for 2 hours at room temperature with an HRP-conjugated rabbit anti-mouse secondary antibody (Agilent, #P0260) diluted 1:2,000. Detection was performed using enhanced chemiluminescent substrate (ThermoFisher Scientific, #34580), and signals were visualized using a Fusion FX7 imaging system (Vilber-Lourmat, Germany). Post-acquisition image adjustments for brightness and contrast were made using ImageJ software retaining relative signal strength.

### Thermal stability of AAV capsids

Thermal stability of the AAV capsids was assessed using the Prometheus NT.48 system (NanoTemper Technologies) with dedicated capillaries (Prometheus NT.48 Capillaries, NanoTemper, PR-C002). Capsid samples were prepared at a concentration of 1 × 10^10^ vg/µL in PBS and loaded in triplicate. Measurements were conducted following the manufacturer’s guidelines, using full excitation intensity and a temperature ramp of 1.5°C per minute from 30°C to 95°C. The melting temperature (Tm) was calculated as the peak of the protein unfolding curve using PR.ThermControl software (version 2.1.6).

### Isolation of PBMCs and activation of T cells

Human peripheral blood mononuclear cells (PBMCs) were isolated from buffy coats of healthy anonymous donors through the DRK Blutspendedienst (Frankfurt, Germany), following ethical approval by the Frankfurt University Hospital ethics committee. PBMCs were isolated via density gradient centrifugation using Histopaque-1077 (Sigma-Aldrich, 10771) to separate them from red blood cells and plasma. After isolation, cells were cryopreserved in a freezing solution composed of 90% fetal bovine serum (FBS) and 10% dimethyl sulfoxide (DMSO). For experimental use, PBMCs were rapidly thawed at 37°C, washed, and cultured in NutriT medium (4Cell Nutri-T; Sartorius, 05-11F2001) containing 0.5% penicillin/streptomycin. T cell activation was induced by plating the cells onto 6-well plates pre-coated with anti-CD3 antibodies (1 µg/ml, Miltenyi Biotec, 130-093-387) and supplementing the medium with soluble anti-CD28 antibodies (3 µg/ml, Miltenyi Biotec, 130-093-375), interleukin-7 (IL-7; 25 U/ml, Miltenyi Biotec, 130-095-361), and interleukin-15 (IL-15; 50 U/ml, Miltenyi Biotec, 130-095-764). On day 3 post-activation, the culture medium was refreshed, maintaining IL-7 and IL-15 supplementation for continued support of T cell proliferation and viability. For ex vivo transduction, activated PBMCs were plated in 96-well plates at a density of 4 × 10⁴ cells per well in 90 µL of NutriT medium supplemented with IL-7 and IL-15. AAV vector stocks were diluted in cytokine-supplemented NutriT medium and added to the cells in a 10 µL volume per well, resulting in a total volume of 100 µL. The following day, the culture medium was replaced with fresh NutriT medium containing IL-7 and IL-15.

### Immunostaining and flow cytometry

Flow cytometric staining was conducted with slight modifications to previously described protocols^19, 22^. Briefly, cells were washed with 1,000 µL of staining buffer (PBS without Ca²⁺/Mg²⁺, supplemented with 2% FBS and 0.1% sodium azide), followed by resuspension in 80–150 µL of the same buffer. Antibody mixes and the viability dye eFluor780 (1:1000 dilution; eBioscience, 65-0865-18) were added to the cells and incubated accordingly. After staining, cells were washed one final time with 1,000 µL of staining buffer. For fixation, cells were subsequently treated with 0.5% paraformaldehyde in PBS lacking divalent cations. Data acquisition was performed on a MACSQuant 10 flow cytometer (Miltenyi Biotec).

### *In vivo* gene transfer

All animal experiments were performed in accordance with the regulations of the German animal protection law and the respective European Union guidelines. Initial biodistribution analysis of luciferase expression was conducted using a subcutaneous tumor model. Crl:NU(NCr)-Foxn1^nu^ mice were subcutaneously engrafted in the left flank with 7 × 10^6^ SKOV3 tumor cells. For orthotopic glioblastoma studies, tumor cell implantation was performed as previously described^29^. Briefly, anesthetized mice were secured in a stereotaxic frame (Stoelting, Wood Dale, USA), and 5 × 10³ GL261-HER2 tumor cells suspended in 2 μL PBS were injected into the right striatum using a 10 μL Hamilton syringe (Hamilton, Bonaduz, Switzerland) and a Quintessential Stereotaxic Injector (Stoelting, Wood Dale, USA). The injection was performed through a burr hole in the skull at a depth of 3 mm from the surface, delivered at a rate of 0.5 μL/min. Following injection, the needle was retracted at a rate of 1 mm/min. Tumor development, body weight, and general health were monitored regularly. In the orthotopic model, successful tumor engraftment was verified by magnetic resonance imaging (MRI).

### Statistics and graphing

All statistical evaluations were conducted using GraphPad Prism (version 9.5.0). The specific statistical tests applied are detailed in the corresponding figure legends. Flow cytometry data were processed and analyzed with FCS Express (version 6.06.0040; De Novo Software) or FlowJo 10.10.0 software (Tree Star Inc), assuming a normal distribution for all comparisons. AlphaFold 3 structural predictions were run on the AlphaFold server ^40^ to de novo fold DARPin displaying AAV9-VP3. AAV9 capsids were generated using the AAV9-VP3 structure (PDB: 3UX1). Resulting structures were visualized with Chimera 1.16 ^41^. Illustrative schematics and graphical elements were generated in part using BioRender.com.

## Data and code availability

All other data generated or analyzed during this study are available from the corresponding author upon reasonable request.

## Acknowledgments

The authors gratefully acknowledge Gundula Braun, Manuela Gallet, and Julia Brynza (Paul-Ehrlich-Institut) for their excellent technical assistance in cloning, AAV vector production, vector genome quantification and western blot.

This work was funded by the BMBF project COMMUTE (16GW0339) awarded to C.J.B.

## Author contributions

Conceptualization: M.B.D., F.S., L.J.Z, F.B.T. and C.J.B. Methodology: M.B.D., F.S., L.J.Z., P.E., F.J., F.B.T. Investigation M.B.D., F.S., L.J.Z, P.E., F.J., and F.B.T. Resources: C.J.B. Writing – original draft: M.B.D. Writing – review & editing: all. Visualization: M.B.D and F.J. Supervision: C.J.B. Project administration: C.J.B. Funding acquisition: T.O. and C.J.B.

---

**Supplementary Fig. 1.**
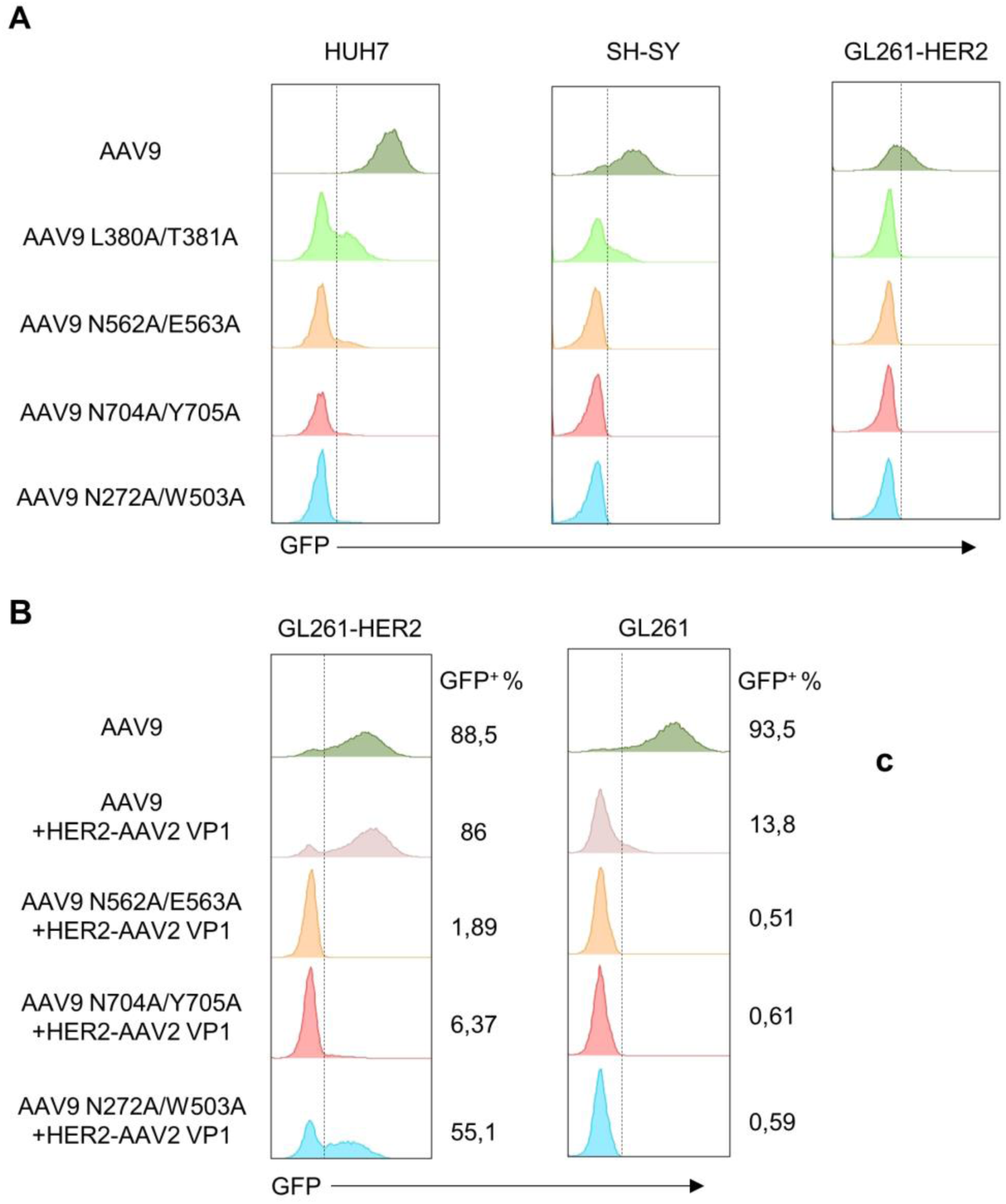
Disruption of natural receptor binding of AAV9 with different blinding mutations and insertion of HER2 DARPin. Transduction of cell lines with AAV9 vectors with different blinding mutations. Vectors are packaging GFP as reporter gene. Representative histograms are shown (n=3). **A)** Comparison in transduction efficiency of unmodified AAV9 versus AAV9 versions with blinding mutations at the position L380A/T381A or N562A/E563A or N704A/Y705A or N272A/W503A. **B)** Comparison in transduction efficiency of unmodified AAV9 versus AAV9 versions with blinding mutations mixed with 1:1 ratio HER2 DARPin-inserted VP1 AAV2 for the production.

**Supplementary Fig. 2.**
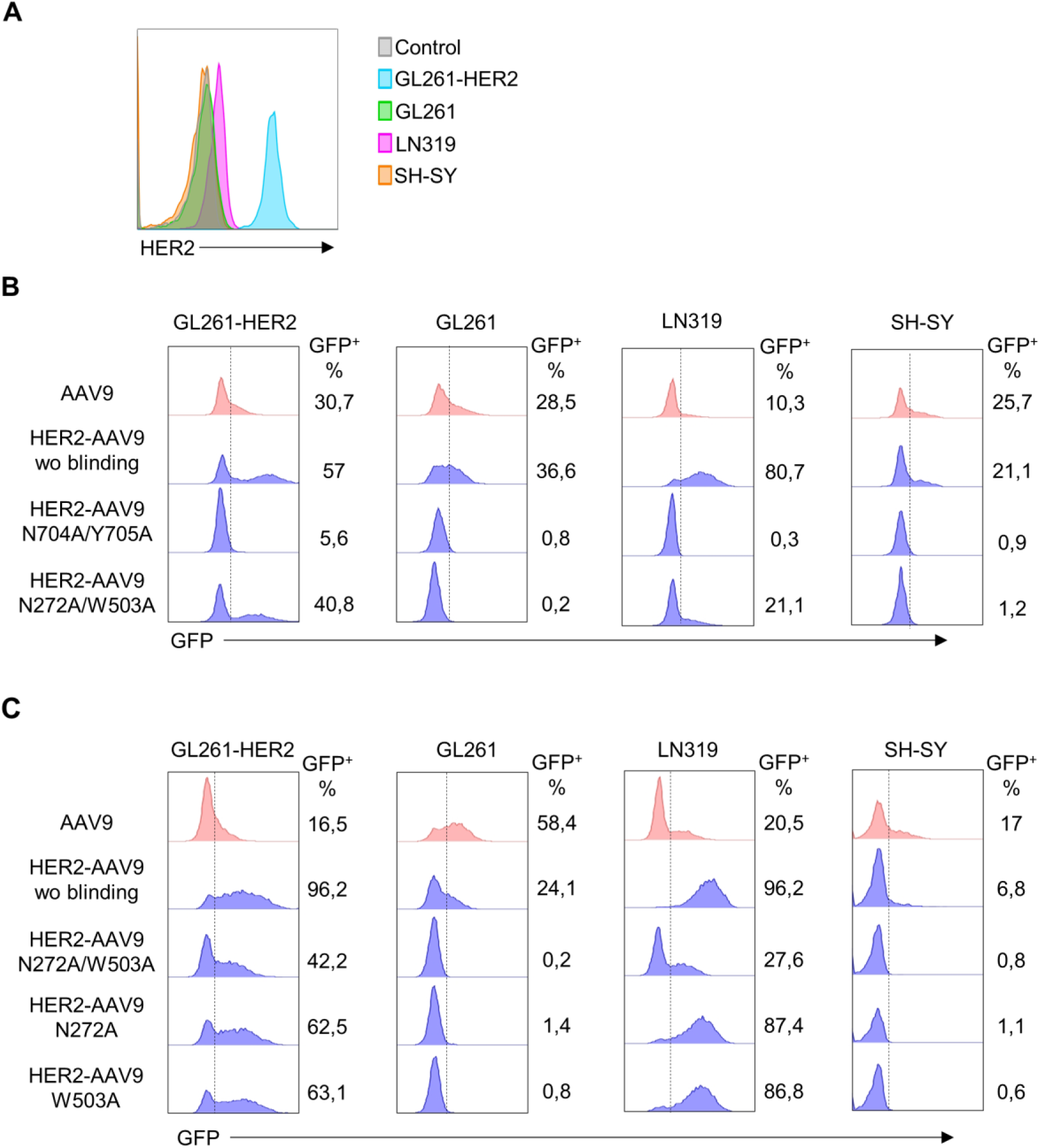
Comparison of HER2 targeted AAV9 versions with different blinding mutations. **A)** Flow cytometry analysis of HER2 expression in HER2 positive GL261-HER2, LN319) and negative (GL261, SH-SY) cell lines. (**B-C**), Transduction of cell lines with HER2 targeted AAV9 vectors with different blinding mutations. Vectors are packaging GFP as reporter gene. Representative fluorescence histograms are shown (n=3). **B)** Comparison in transduction efficiency of unmodified AAV9 versus HER2-AAV9 wo blinding, HER2-AAV9 N704A/Y705A or HER2-AAV9 N272A/W503A. **C)** Comparison in transduction efficiency of unmodified AAV9 versus HER2-AAV9 wo blinding, HER2-AAV9 N272A/W503A, HER2-AAV9 N272A or HER2-AAV9 W503A. Representative fluorescence histograms are shown (n=3).

**Supplementary Fig. 3.**
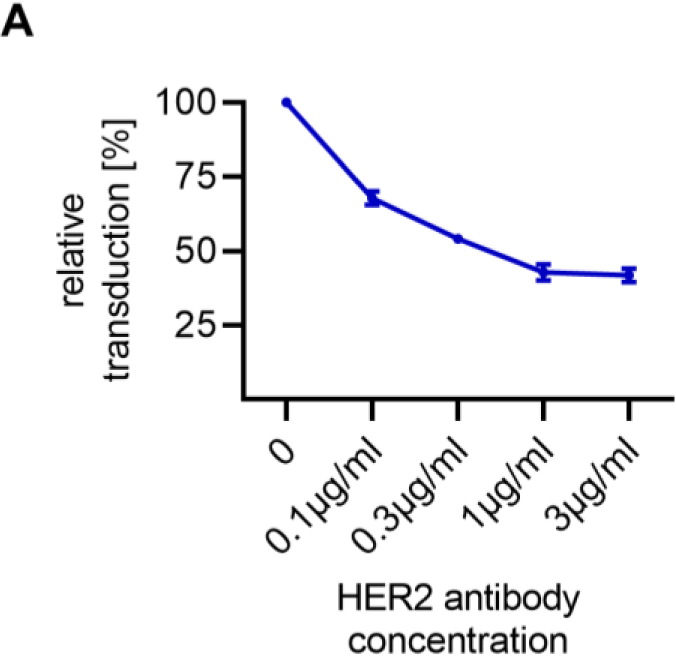
Supplementary data to Fig. 1. Transduction of SKOV3 cells with HER2-AAV9 vectors with presence of increasing concentrations of anti-HER2 antibody. Transduction efficiency was quantified by flow cytometry, and relative transduction was calculated based on mean fluorescence intensity (MFI) values (n=3).

**Supplementary Fig. 4.**
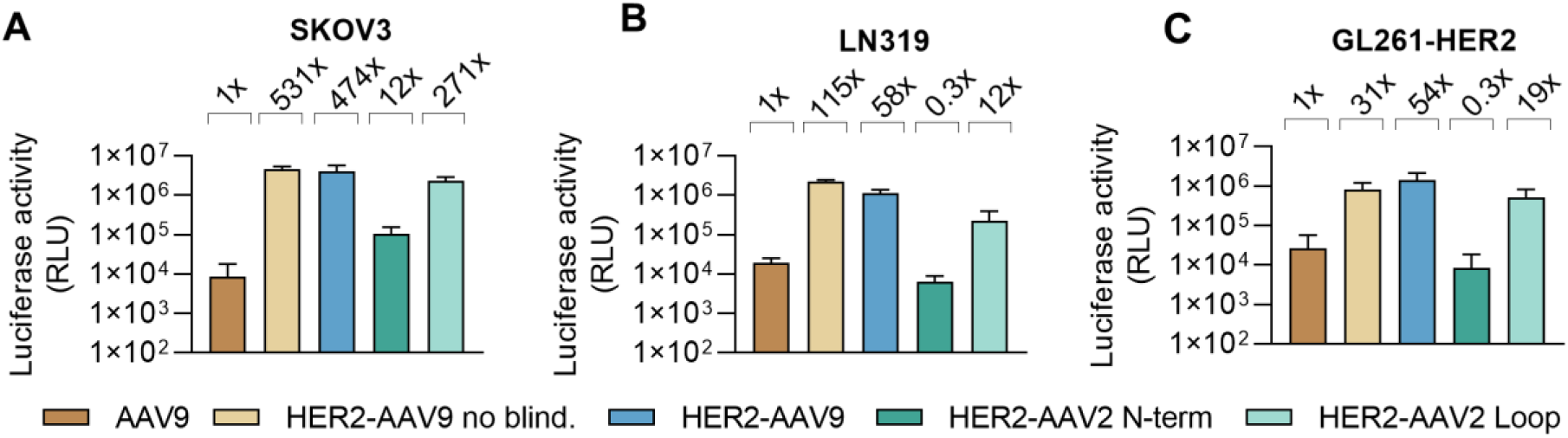
Complementary *in vitro* data to Fig. 2. A-C) *In vitro* transduction on cell lines. Luciferase reporter gene transfer was assessed using the indicated AAV vectors in HER2-positive cells SKOV3 **(A)**, LN319 **(B)**, and GL261-HER2 **(C)**. Cells were incubated with each vector carrying luciferase transgene at 1x10^5^ vg/cell, and luciferase expression was subsequently measured to evaluate transduction efficiency (n=3). The fold increase in transduction relative to AAV9 is shown above each bar. Bars represent means, error bars represent SD.

**Supplementary Fig. 5.**
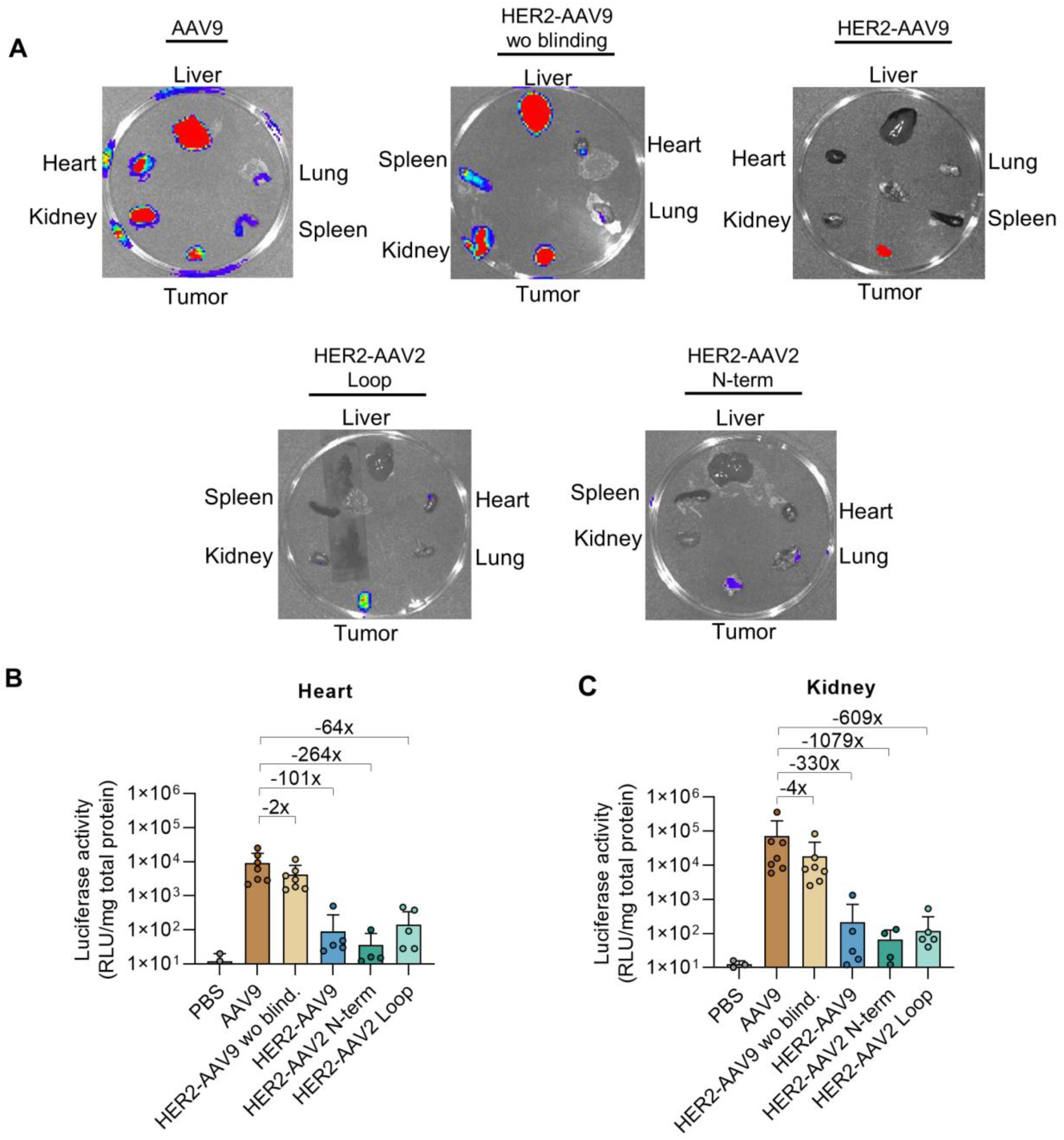
Complementary data to Fig. 2. *In vivo* transduction in a subcutaneous tumor model. Nude mice were engrafted with SKOV3 cells in the left flank and systemically injected via the tail vein with each luciferase-encoding AAV vectors (2x10^11^ vg). **A)** Representative IVIS images of isolated organs at day 7 post-injection are shown. Following euthanasia, tumors and organs were harvested, lysed in luciferase lysis buffer, and analyzed to determine transduction efficiency in hearth **(B)** and kidney **(C)**. Each dot represents an individual mouse. The fold increase in transduction relative to AAV9 is indicated above each bar. Bars represent mean values; error bars represent SD.

**Supplementary Fig. 6.**
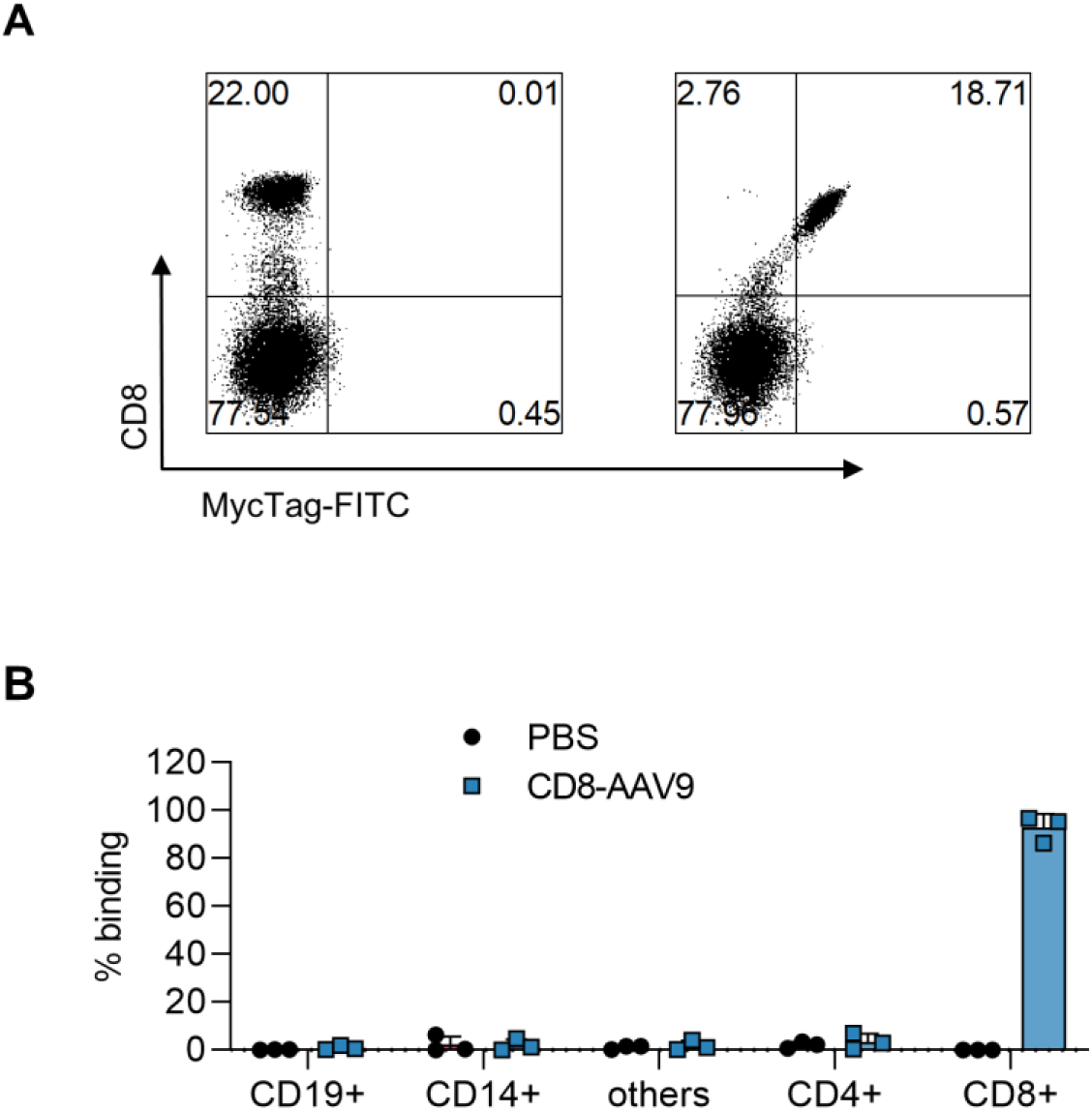
CD8-AAV9 provides specific binding to CD8+ T cells. Binding of CD8-AAV9 to human PBMCs. Peripheral blood from three donors were incubated with the indicated AAVs at 1 × 10^11^ vg/cell or PBS for 4 h at 37°C. Percentages of AAV-bound cells were determined by flow cytometry after staining with the Myc-tag antibody.

**Table S1:** Batch-to-batch variability of various AAV types.

| Type of AAV | Batch | Packaged gene | Genome copies [gc/ml] |
| --- | --- | --- | --- |
| AAV9 | #1 | luciferase-GFP | 6.8E+13 |
|  | #2 | GFP | 8.1E+13 |
|  | #3 | luciferase-GFP | 8.9E+13 |
| HER2-AAV9<sup>noblind</sup> | #1 | luciferase-GFP | 6.5E+13 |
|  | #2 | GFP | 4.3E+14 |
|  | #3 | luciferase-GFP | 1.2E+14 |
| HER2-AAV9 | #1 | luciferase-GFP | 5.6E+13 |
|  | #2 | GFP | 5.1E+14 |
|  | #3 | luciferase-GFP | 1.1E+14 |

